# Efficiency of local learning rules in threshold-linear associative networks

**DOI:** 10.1101/2020.07.28.225318

**Authors:** Francesca Schönsberg, Yasser Roudi, Alessandro Treves

## Abstract

We show that associative networks of threshold linear units endowed with Hebbian learning can operate closer to the Gardner optimal storage capacity than their binary counterparts and even surpass this bound. This is largely achieved through a sparsification of the retrieved patterns, which we analyze for theoretical and empirical distributions of activity. As reaching the optimal capacity via non-local learning rules like back-propagation requires slow and neurally implausible training procedures, our results indicate that one-shot self-organized Hebbian learning can be just as efficient.

## INTRODUCTION

Local learning rules, those that change synaptic weights depending solely on pre- and post-synaptic activation, are generally considered to be more biologically plausible than non-local ones. They can be implemented through extensively studied processes in the synapses [1] and they allow neural networks to self-organise into content-addressable memory devices [2–4]. But how effective are local learning rules? Quite ineffective, has been the received wisdom since the 80’s, when back-propagation algorithms came to the fore. However, this common sense is based on analysing networks of binary units [3, 5–7], while neurons in the brain are not binary.

A better, but still mathematically simple description of neuronal input-current-to-impulse-frequency transduction is via a threshold-linear transfer function [8–10], which also represents the activation function predominantely adopted in recent deep learning applications [11–14]) (often called, in that context, Rectified Linear units or ReLu). Therefore, one may ask whether the results from the 80’s highlighting the contrast between the effective, iterative procedures used in machine learning and the self-organized, one-shot, perhaps computationally ineffective Hebbian learning are valid beyond binary units [15].

The Hopfield model [3], which includes a simple *Hebbian* [2] prescription for structuring all the connection weights in one go, had been analysed and found to be able to retrieve only up to *p_max_* ≃ 0.14N activity patterns, in a network of *N* binary units [5], with *C* = *N* – 1 input connections per unit. In contrast, Elizabeth Gardner showed [7] that the optimal capacity such a network can attain is *p_max_* = 2*C*, about 14 times higher. This optimal capacity can be approached with iterative procedures based on the notion of a desired output for each unit, and of progressively reducing the difference between current and desired output – like backpropagation, in multi-layer neural networks. This consolidated the impression that unsupervised, Hebbian plasticity may well be of biological interest, but is rather inefficient and unsuitable for performance-driven machine learning applications. The negative characterization was not redeemed by the finding that in sparsely connected nets, those where *C* ≪ *N*, the pattern capacity *α_c_* = *p_max_/C* can be closer to the Gardner bound (a factor 3 away) [6]; and approach it, when the coding is sparse, i.e., the fraction of units active in each pattern is *f* ≪ 1 [16]. What about TL units? Are they more efficient in the unsupervised learning of memory patterns?

Here we derive and evaluate a closed set of equations for the optimal pattern capacity *à la Gardner* in networks of TL units, as a function of the fraction *f* of active units, and test our results by learning those weights with a TL perceptron. We show that first, while for stored patterns that are binary, such errorless capacity is larger than the Hebbian capacity no matter how sparse the code, this does not, in general, hold for non-binary stored patterns, and for other distributions the Hebbian capacity can even surpass the Gardner bound. This perhaps surprising violation of the bound is because the Gardner calculation imposes an infinite output precision [17], while Hebbian learning exploits its loose precision to *sparsify* the retrieved pattern. In other words, with TL units, Hebbian capacity can get much closer to the optimal capacity or even surpass it, by permitting errors and retrieving a sparser version of the stored pattern. Experimentally observed firing activity distributions from the inferior-temporal cortex [18], which can be taken as patterns to be stored, would be sparsified about 50% by Hebbian learning, and would reach about 50% – 80% of the Gardner capacity.

## MODEL DESCRIPTION

We consider a network of *N* units and *p* patterns of activity, 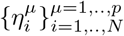 each representing one memory stored in the connection weights via some procedure. Each 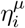 is drawn independently for each unit *i* and each memory *μ* from a common distribution Pr(*η*). We denote the activity of each unit *i* by *v_i_* and assume that it is determined by the activity of the *C* units feeding to it, through a TL activation function

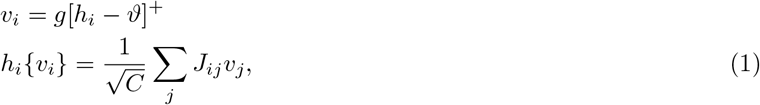

where [*x*]^+^ = *x* for *x* > 0 and = 0 otherwise; and both the gain *g* and threshold *ϑ* are fixed parameters taken to be set for the whole network. In a fully connected, recurrent network, *C* = *N* – 1, but we intend to consider also more general cases of diluted connectivity. The *storage capacity α_c_* ≡ *p_max_*/*C*, is then the maximal number of memories that the network can store and individually retrieve, per unit. The synaptic weights *J_ij_* are taken to satisfy the spherical normalization condition for all *i*

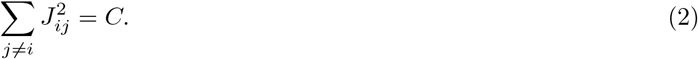

We are interested in finding the set of *J_ij_* that satisfy Eq. (2), such that patterns 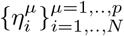 are self-consistent solutions of Eqs. 1, namely that for all *i* and *μ* we have, 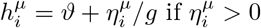 and 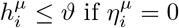.

## REPLICA ANALYSIS

Adapting the procedure introduced by Elisabeth Gardner [7] for binary units to our network, we evaluate the fractional volume of the space of the interactions *J_ij_* which satisfy Eqs. (2) and (1); and using the replica trick we obtain the standard order parameters 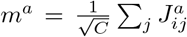 and 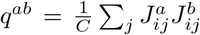 corresponding, respectively, to the average of the weights within each replica and to their overlap between replicas (Supplemental Material at [URL], Sect. A). Assuming the replica symmetric ansatz simplifies *q^ab^* ≡ *q* and *m^a^* ≡ *m*. We focus on increasing the number p of stored memories, in the *C* → ∞ limit, up to when the volume of the compatible weights shrinks to a single point, i.e., there is a unique solution, and the maximal storage capacity has been reached. For this purpose we take the limit *q* → 1, corresponding to the case where all the replicated weights are equal, implying that only one configuration exist which satisfies the equations. Adding a further memory pattern would make it impossible, in general, to satisfy all equations.

We have derived a system of two equations for the maximal storage capacity *α_c_* = *p*_max_/*C* for a network of threshold-linear units

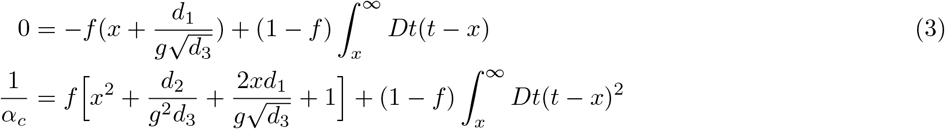

where we have introduced the averages over 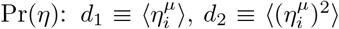 and 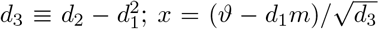 is the normalized difference between the threshold and the mean input, while *f* = Pr(*η* > 0) is the fraction of active units. The two equations yield *x* (hence implicitly setting an optimal value for *ϑ*) and *α_c_*. Note that both equations can be understood as averages over units, respectively of the actual input and of the square input, which determine the amount of quenched noise and hence the storage capacity.

The storage capacity then depends on the proportion *f* of active units, but also on the gain *g*, and on the moments of the distribution Pr(*η*), di and *d*_3_. As can be seen in Fig. 1a, at fixed *g*, the storage capacity increases as more and more units remain below threshold, ceasing to contribute to the quenched noise. In fact, the storage capacity diverges, for *f* → 0, as

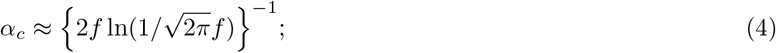

see Supplemental Material at [URL], Sect. B. for the full derivation. At fixed *f*, there is an initially fast increase with *g* followed by a plateau dependence for larger values of *g*. One can show that 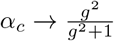 as *f* → 1, i.e., when all the units in the memory patterns are above threshold, it is always *α_c_* < 1 for any finite *g*. At first sight this may seem absurd: a linear system of *N*^2^ independent equations and *N*^2^ variables always has an inverse solution, which would lead to a storage capacity of (at least) one. Similar to what already noted in [17], however, the inverse solution does not generally satisfy the spherical constraint in Eq. (2); but it does, in our case, in the limit *g* → ∞ and this can also be understood as the reason why the capacity is highest when *g* is very large. In practice, Fig. 1 indicates that over a broad range of f values the storage capacity approaches its *g* → ∞ limit already for moderate values of the gain; while the dependence on *d*_1_ and *d*_3_ is only noticeable for small *g*, as can be seen by comparing Fig. 1c and d. In the *g* → ∞ limit, one sees that Eqs. (3) depend on Pr(*η*) only through *f*.

**FIG. 1.**
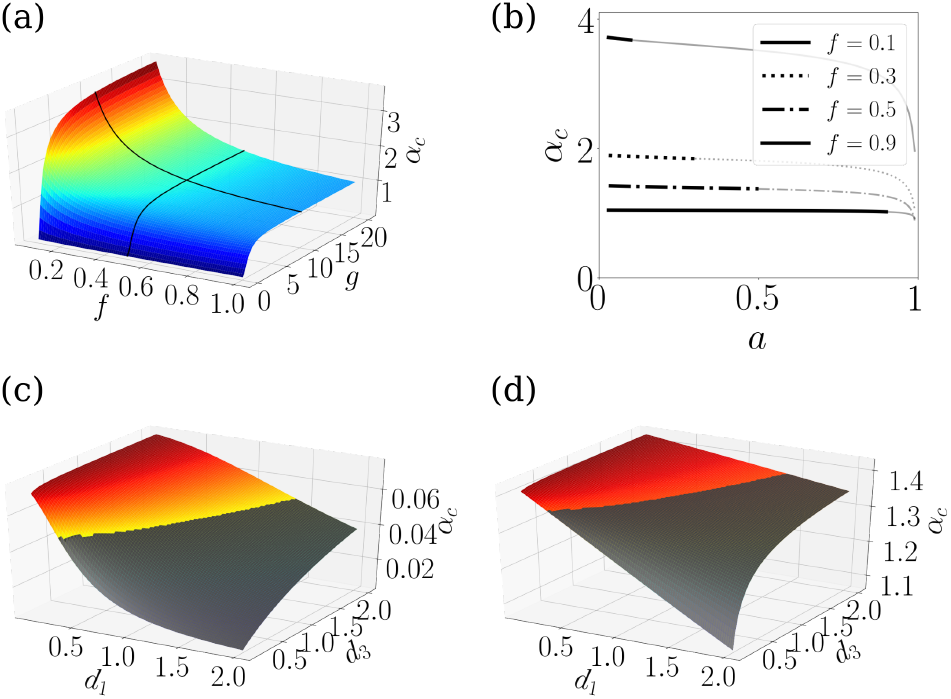
Dependence of the Gardner capacity *α_c_* on different parameters. *α_c_* plotted in (a) as a function of *g* and *f*(*d*_1_ = 1.1, *d*_2_ = 2), in (b) as a function of 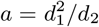 for different values of *f*(*g* = 10, *d*_1_ = 1.1) in (c) and (d) as a function of di and *d*_3_ for *g* = 0.2 and *g* = 10, respectively (*f* = 0.5). Note that fixing *f*, restricts the available range of *a*, as *a* cannot be larger than *f*; the inaccessible ranges of f values are shadowed in (b), (c) and (d).

Eqs. (3), at *g* → ∞, have been verified by explicitly training a threshold linear perceptron of *N* = 100 units and *C* = *N* connections, with *p* binary patterns. We have evaluated *α_c_* = *p_max_*/*C* numerically by estimating *p_max_* as the maximal number of patterns which can be retrieved with no errors. The numerical values for the storage capacity are depicted as red diamonds in Fig. 2, it can be noticed that they follow the profile of the solid line describing the *g* → ∞ limit of Eq. (3). See Supplemental Material at [URL], Sect. C for details about the algorithm.

**FIG. 2.**
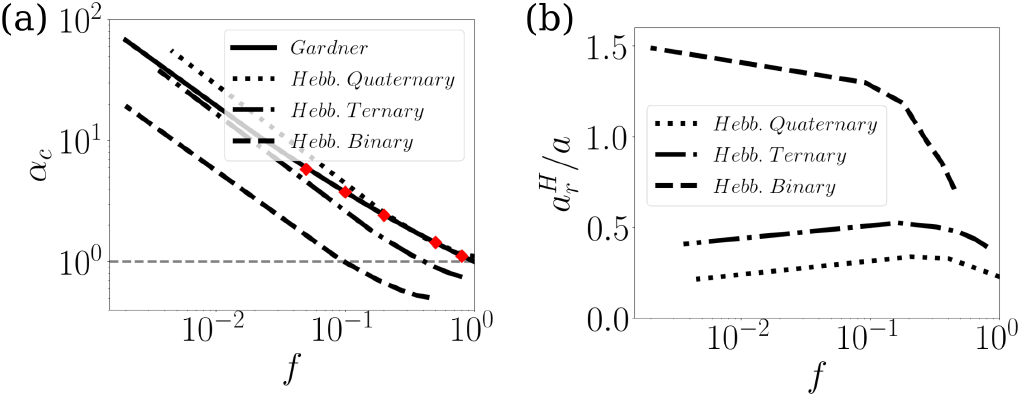
Hebbian capacity vs Gardner bound. (a) 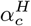 as a function of *f* for different sample distribution of stored patterns compared to the universal 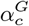 bound for errorless retrieval, i.e. the g → ∞ limit of Eq (3); the red diamonds are reached with explicit TL perceptron training. (b) the sparsification of the stored patterns at retrieval, for Hebbian networks loaded at their capacity.

## COMPARISON WITH A HEBBIAN RULE: THEORETICAL ANALYSIS

As described in the Introduction, when used in a fully connected Hopfield network of binary units, and no sparse coding, Hebbian learning could only reach ~ 1/14 of the Gardner bound. With very sparse connectivity, however, the same network can get to 1/π of the bound, even if it gets there through a second-order phase transition, where the retrieved pattern has vanishing overlap with the stored one [6]. With TL units, the appropriate comparison to the capacity *à la Gardner* is therefore that of a Hebbian network with *extremely diluted* connectivity, as in this limit the weights *J_ij_* and *J_ji_* are effectively independent and noise reverberation through loops is negligible. The capacity of such a sparsely connected TL network was evaluated analytically in [19]. Whereas in the *g* → ∞ limit the Gardner capacity depends on Pr(*η*) only via *f*, for Hebbian networks it does depend on the distribution, and most importantly on, *a*, the *sparsity*

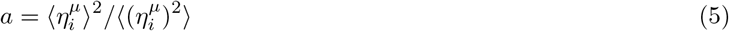

whose relation to *f* depends on the distribution [19].

Here we focus on 3 examples of binary, ternary and quaternary distributions (Supplemental Material at [URL], Sect. D); in these examples, the parameters f and *a* are related through *f* = *a*, 9*a*/5 and 9*a*/4, respectively. The results of the numerical comparisons are shown in Fig. 2. Both the Hebbian and the Gardner capacity diverge in the sparse coding limit.

One can see in Fig. 2a that when attention is restricted to binary patterns, the Gardner capacity *seems* to provide an upper bound to the capacity reached with Hebbian learning; more structured distributions of activity, however, dispel such a false impression: the quaternary example already shows higher capacity for sufficiently sparse patterns.

The bound, in fact, would only apply to perfect errorless retrieval, whereas Hebbian learning creates attractors which are, up to the Hebbian capacity limit, correlated but not identical to the stored patterns, similarly to what occurs with binary units [5]; in particular, we notice that when considering TL units and Hebbian learning, in order to reach close to the capacity limit, the threshold has to be such as to produce sparser pattern at retrieval, in which only the units with the strongest inputs get activated. The sparsity 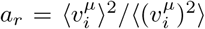 of the retrieved memory can be calculated [19], see Supplemental Material at [URL], Sect. D for an outline. Fig. 2b shows the ratio of the sparsity of the retrieved pattern produced by Hebbian learning, denoted as 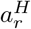, to that of the stored pattern *a*, vs. *f*. As one can see, except for the binary patterns at low *f*, the retrieved patterns, at the storage capacity, are always sparser than the stored ones. The largest sparsification happens for quaternary patterns, for which the Hebbian capacity overtakes the bound on errorless retrieval, at low *f*. Sparser patterns emerge as, to reach close to 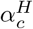, *ϑ* has to be such as to inactivate most of the units with intermediate activity levels in the stored pattern. Of course, the perspective is different if 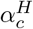 is considered as a function of *a_r_* instead of *a*, in which case the Gardner capacity remains unchanged, as it implies retrieval with *a_r_* = *a*, and above 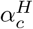 for each of the 3 sample distributions; see Fig. 1 of Supplemental Material at [URL].

## COMPARISON WITH A HEBBIAN RULE: EXPERIMENTAL DATA

Having established that the Hebbian capacity of TL networks can surpass the Gardner bound on errorless retrieval for some distributions, we ask what would happen with the distribution of firing rates naturally occurring in the brain. Here we consider published distributions of single units in infero-temporal visual cortex, while the animals were watching short naturalistic movies [18]. Such distributions can be taken as examples of patterns elicited by the visual stimulus, and to be stored with Hebbian learning, given appropriate conditions, and later retrieved using attractor dynamics, triggered by a partial cue [20–24]. How many such patterns can be stored, and with what accompanying sparsification?

Fig. 3a and b show the analysis of two sample distributions (of the top right and top left cells in Fig. 2 of [18]). The observed distributions, in blue, with the “Gardner” label, are those we assume could be stored, and which could be retrieved exactly as they are with a suitable training procedure bound by the Gardner capacity. In orange, instead, we plot the distribution that would be retrieved following Hebbian learning operating at its capacity, see Supplemental Material at [URL], Sect. F., for the estimation of the retrieved distribution. Note that the absolute scale of the retrieved firing rate is arbitrary, what is fixed is only the shape of the distribution, which is sparser (as clear already from the higher bar at zero). The pattern in Fig. 3a, which has *a* < 0.5, could also be fitted with a one-parameter exponential shape having *f* = 2*a* (see Supplemental Material at [URL], Sect. E). In that panel we also report the values of the 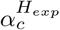 and 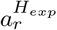 calculated assuming the continuous exponential instead of the observed discrete distribution (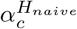 and 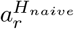). Fig. 3c shows both 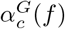 and 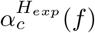; on top of these curves and in the inset we have indicated as diamonds the values calculated for the 9 empirical distributions present in [18] and as circles the fitted values for those which could be fitted to an exponential.

**FIG. 3.**
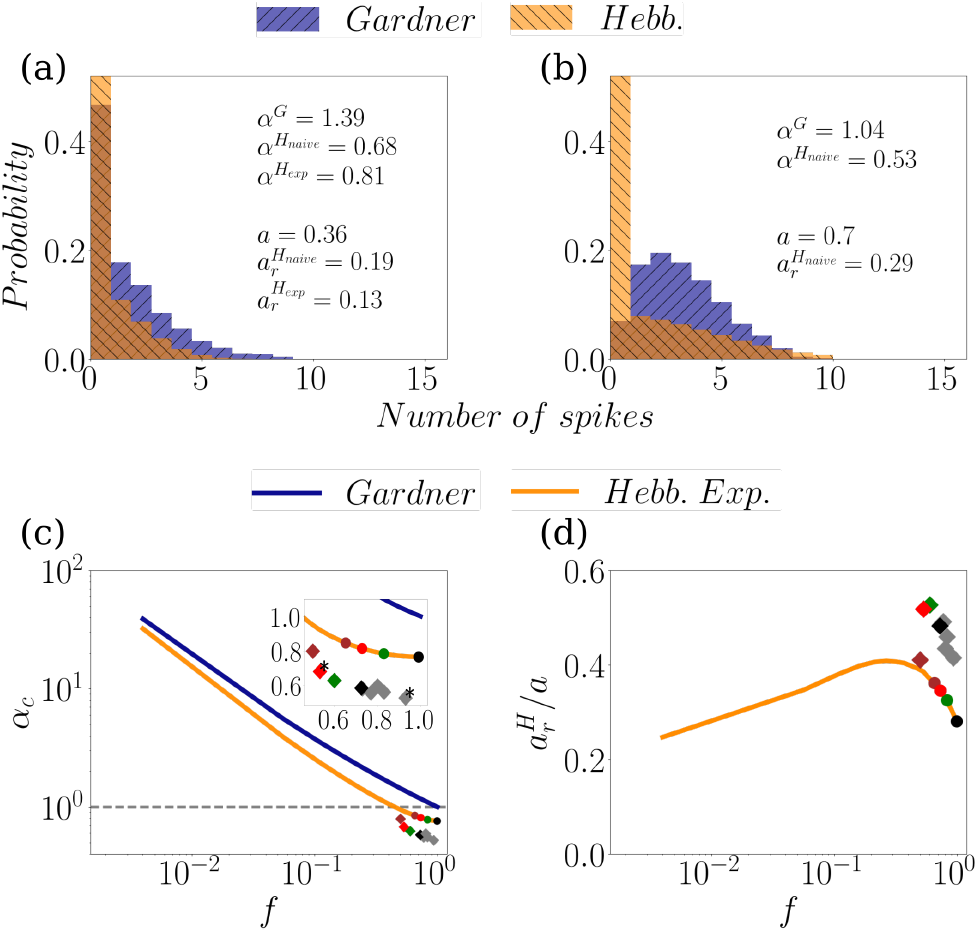
Hebbian learning vs. the Gardner errorless bound for experimental data. (a,b) Examples of the histograms of two experimentally recorded spike counts (blue) and the retrieved distribution, if the patterns were stored using Hebbian learning (orange). Note that the retrieved distributions *à la Gardner* would be the same as the stored patterns. (c) Analytically calculated universal Gardner capacity 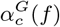 (blue), i.e. the *g* → ∞ limit of Eq (3), compared to 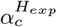 for the Hebbian learning of an exponential distribution (orange). The diamonds are the values 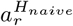 achieved with the 9 original discrete distributions, and the circles the values 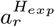 for those 4 that can be fit to an exponential distribution. The asterisk marks the two cells whose distribution is plotted in a) and b). d) Sparsification of the retrieved patterns, for Hebbian learning.

There are three conclusions that we can draw from these data. First, the Hebbian capacity from the empirical distributions is about 80% of that of the exponential fit, when available. Second, for distributions like those of these neurons, the capacity achieved by Hebbian learning is about 50% – 80% of the Gardner capacity for errorless retrieval, depending on the neuron and whether we take its discrete distribution “as is”, or fit it with a continuous exponential. Third, Hebbian learning leads to retrieved patterns which are 2 – 3 times sparser than the stored patterns, again depending on the particular distribution and whether we take the empirical distributions or their exponential fit. The empirical distributions achieve a lower capacity than that of their exponential fit, which leads to further sparsification at retrieval. This is illustrated in Fig. 3d, which shows the ratio of the sparsity of patterns retrieved after Hebbian storage to that of the originally stored pattern, vs. *f*.

## DISCUSSION

The general notion of attractor neural networks (ANN) has been instrumental in conceptualizing the storage of long-term memories, including those experimentally accessible, such as spatially selective memories in rodents. For example, the activity of Place [25, 26] and Grids cells [27] has been analyzed in terms of putative low-dimensional attractor manifolds [28, 29]. In detail, however, the applicability of advanced mathematical results [30, 31] has been challenged by several cortically implausible assumptions incorporated in the early models. In particular, an understanding of the effectiveness of Hebbian learning has been hampered by the fact that a natural benchmark, the Gardner bound on the storage capacity, had been derived initially only for binary units. The bound for binary units, as described in the Introduction, was found to be far above the Hebbian capacity, but then no binary or quasi-binary pattern of activity has ever been observed in the cerebral cortex. A few studies have considered non-binary units: TL networks have been shown to be less susceptible to spin-glass effects [32] and to mix-ups of memory states [33] but in the framework of *à la Gardner* calculations they have focused on other issues than associative networks storing sparse representations. For instance in [17] the authors carried out a replica analysis on a generic gain function, but then focused their study on the tanh activation function and on activity patterns that were not neurally motivated. Clopath and Brunel [34] considered monotonically increasing activation functions under the constraint of non-negative weights as a model of cerebellar Purkinje cells and, more recently, it has been shown with a replica analysis [35] that TL units can largely tolerate perturbations in the exact values of the weights and of the inputs.

Here, we report the analytical derivation of the Gardner capacity for networks of TL units, validate the result with TL perceptron training, and compare it with the performance of networks with Hebbian weights. We argue that the comparison has to be framed in the context of the difference between stored and retrieved patterns, that becomes more salient, the higher the storage load of the Hebbian network. It remains to be assessed whether a stability parameter, comparable to the *κ* used in the original Gardner calculations [7], could be considered also for TL units. Further understanding would also derive from the comparison between the maximal information content per synapse when patterns are stored via Hebbian or iterative learning, as previously performed for binary units [36].

For typical cortical distributions of activity in visual cortex, Hebbian one-shot local learning leads to utilize already 50% – 80% of the available capacity for errorless retrieval, leading to markedly sparser retrieved activity. In the extreme in which only the most active cells remain active, those retrieved memories cannot be regarded as the full pattern, with its entire information content, but more as a pointer, effective perhaps to address the full memory elsewhere, as posited in *index* theories of 2-stage memory retrieval [37].

## ACKNOWLEDGMENTS

We thank Rémi Monasson and Aldo Battista for useful comments and discussions. This research received funding from the EU Marie Skłodowska-Curie Training Network 765549 “M-Gate”, and partial support from Human Frontier Science Program RGP0057/2016, the Research Council of Norway (Centre for Neural Computation, grant number 223262; NORBRAIN1, grant number 197467), and the Kavli Foundation.

