## Supplemental Material for "Efficiency of local learning rules in threshold-linear associative networks"

#### A. Derivation of the storage capacity

We start by considering a single threshold-linear unit whose activity is denoted by $u$. The neuron receives $C$ inputs $v_j$, for $j = 1 \cdots C$ through synaptic weights $J_j$. The activity of the neuron is determined through the threshold-linear activation function as

$$u = g[h_i - \vartheta]^+$$
$$h\{v\} = \frac{1}{\sqrt{C}} \sum_j J_j v_j, \tag{1}$$

We assume that we have $p$ patterns, indexed as $\mu = 1 \cdots p$, of activity over the inputs that we denote by $\xi_j^\mu$. To each input pattern $\mu$ we also consider a desired output activity by the neuron that we denote $\eta^\mu$. We are interested in finding how many patterns can be stored in the synaptic weights, such that the input activity elicits the desired output activity, assuming that the synaptic weights satisfy the spherical constraint

$$\sum_{j \neq i} J_j^2 = C. \tag{2}$$

This task, essentially boils down to calculating the expectation of the logarithm of the fractional volume $V$ of the interaction space over the distribution of $\eta$ and $\xi$, defined as

$$V = \frac{\int \prod_j dJ_j \delta\left(\sum_j J_j^2 - C\right) \prod_\mu \left[(1-\delta_{\eta^\mu,0})\,\delta\left(h^\mu - \vartheta - \frac{\eta^\mu}{g}\right) + \delta_{\eta^\mu,0}\Theta\left(\vartheta - h_i^\mu\right)\right]}{\int \prod_j dJ_j \prod_i \delta\left(\sum_j J_j^2 - C\right)}, \tag{3}$$

For calculating $\langle \log V \rangle_{\eta,\xi}$, we use the replica trick. The initial task is to compute the replicated average $\langle V^n \rangle_\xi$, namely

$$\langle V^n \rangle_\xi = \prod_{a=1,..,n} \prod_\mu \frac{\int \prod_j dJ_j^a \delta\left(\sum_j (J_j^a)^2 - C\right) \left\langle (1-\delta_{\eta^{a,\mu},0})\,\delta\left(h^{a,\mu} - \vartheta - \frac{\eta^\mu}{g}\right) + \delta_{\eta^{a,\mu},0}\Theta(\vartheta - h^{a,\mu})\right\rangle}{\int \prod_{j,j\neq i} dJ_j^a \delta\left(\sum_j (J_j^a)^2 - C\right)}. \tag{4}$$

We first compute the numerator. To compute the averages over $\xi$ in the numerator, we note that the delta function can be written as

$$\begin{aligned}\delta(h^{a,\mu} - \vartheta - \frac{\eta^\mu}{g}) &= \int \frac{dx_\mu^a}{2\pi} \exp\left\{ix_\mu^a\Big(\frac{1}{\sqrt{C}}\sum_j J_j^a \eta^\mu - \vartheta - \frac{\eta^\mu}{g}\Big)\right\} \\ &= \int \frac{dx_\mu^a}{2\pi} \exp\Big[-\frac{ix_\mu^a}{g}\Big(\eta^\mu + g\vartheta\Big)\Big] \exp\Big[\frac{ix_\mu^a \sum_j J_j^a \eta^\mu}{\sqrt{C}}\Big].\end{aligned} \tag{5}$$

For the average of the Heaviside function, we write

$$\begin{aligned}\Theta(\vartheta - h^{a,\mu}) &= \int_0^\infty d\lambda_\mu^a \delta[\lambda_\mu^a - (\vartheta - h^{a,\mu})] \\ &= \int_0^\infty \frac{d\lambda_\mu^a}{2\pi} \int_{-\infty}^\infty dy_\mu^a \exp[iy_\mu^a(\lambda_\mu^a - (\vartheta - h^{a,\mu}))] \\ &= \int_0^\infty \frac{d\lambda_\mu^a}{2\pi} \int_{-\infty}^\infty dy_\mu^a \exp\Big[iy_\mu^a(\lambda_\mu^a - \vartheta)\Big] \exp\Big[\frac{iy_\mu^a \sum_j J_j^a \eta^a}{\sqrt{C}}\Big].\end{aligned} \tag{6}$$

We now use the above identities in Eqs. (5) and (6) to compute the following quantity that appears in the numerator of Eq. (4), assuming independently drawn $\xi$ as

$$\begin{aligned}e^{CM} &\equiv \left\langle \prod_{\mu,a} (1-\delta_{\eta^{a,\mu},0})\delta(h^{a,\mu} - \vartheta - \frac{\eta^\mu}{g}) + \delta_{\eta^{a,\mu},0}\Theta(\vartheta - h^{a,\mu}) \right\rangle_{\xi,\eta} \\ &= \prod_\mu \left\langle (1-\delta_{\eta^{a,\mu},0}) \Big\langle \prod_a \delta(h^{a,\mu} - \vartheta - \frac{\eta^\mu}{g}) \Big\rangle_{\xi^\mu} + \delta_{\eta^{a,\mu},0} \Big\langle \prod_a \Theta(\vartheta - h^{a,\mu}) \Big\rangle_{\xi^\mu} \right\rangle_{\eta^\mu}.\end{aligned} \tag{7}$$

In order to compute the average of the *delta* functions in Eq.(7), we use the approximation

$$\langle \exp(x) \rangle \approx \exp\left\{ \langle x \rangle + \frac{\langle x^2 \rangle}{2} - \frac{\langle x \rangle^2}{2} \right\} \tag{8}$$

to calculate the following average

$$\begin{aligned}
&\Big\langle \exp\left\{ \frac{i \sum_{a,j} x_\mu^a J_j^a \xi_j^\mu}{\sqrt{C}} \right\} \Big\rangle_{\xi^\mu} = \\
&= \exp\left\{ \frac{i}{\sqrt{C}} \sum_{a,j} x_\mu^a J_j^a \langle \xi_j^\mu \rangle - \frac{1}{2C} \sum_{a,b,j,k} x_\mu^a x_\mu^b J_j^a J_k^b \langle \xi_j^\mu \xi_k^\mu \rangle - \frac{1}{2} \left( \frac{i}{\sqrt{C}} \sum_{a,j} x_\mu^a J_j^a \langle \xi_j^\mu \rangle \right) \left( \frac{i}{\sqrt{C}} \sum_{b,k} x_\mu^b J_j^b \langle \xi_k^\mu \rangle \right) \right\} \\
&= \exp\left\{ \frac{i}{\sqrt{C}} \sum_{a,j} x_\mu^a J_j^a \langle \xi_j^\mu \rangle - \frac{1}{2C} \sum_{a,b,j} x_\mu^a x_\mu^b J_j^a J_j^b \langle (\xi_j^\mu)^2 \rangle + \frac{1}{2C} \sum_{a,b,j} x_\mu^a x_\mu^b J_j^a J_j^b \langle \xi_j^\mu \rangle^2 \right\}
\end{aligned} \tag{9}$$

where in going from the second to third line in Eq. (9), we have used the fact that $\langle \xi_j^\mu \xi_k^\mu \rangle = \langle \xi_j^\mu \rangle \langle \xi_k^\mu \rangle$. Expanding the second exponential in the second line of Eq. (5), we can write in the large $C$ limit

$$\begin{aligned}
&\Big\langle \prod_a \delta(h^{a,\mu} - \vartheta - \frac{\eta^\mu}{g}) \Big\rangle_{\xi^\mu} = \\
&= \int_{-\infty}^{\infty} \Big[ \prod_a \frac{dx_\mu^a}{2\pi} \Big] \exp\Big[ -\frac{i}{g}(\eta^\mu + g\vartheta) \sum_a x_\mu^a + i d_1^{inp} \sum_a x_\mu^a m^a - \frac{d_3^{inp}}{2} \Big( \sum_a (x_\mu^a)^2 + 2 \sum_{a<b} x_\mu^a x_\mu^b q^{ab} \Big) \Big] \\
&\equiv I_1(q^{ab}, m^a, \eta^\mu)
\end{aligned} \tag{10}$$

in which we have assumed symmetric replicas and defined $d_1^{inp} \equiv \langle \xi_j^\mu \rangle$, $d_2^{inp} \equiv \langle (\xi_j^\mu)^2 \rangle$, $d_3^{inp} \equiv d_2^{inp} - (d_1^{inp})^2$ and

$$q^{ab} = \tfrac{1}{C} \textstyle\sum_j J_j^a J_j^b \tag{11a}$$

$$m^a = \tfrac{1}{\sqrt{C}} \textstyle\sum_j J_j^a \tag{11b}$$

Similarly, using the identity in Eq. (6) we have

$$\begin{aligned}
&\Big\langle \prod_a \Theta(\vartheta - h^{a,\mu}) \Big\rangle_{\xi^\mu} = \\
&= \int_0^{\infty} \Big[ \prod_a \frac{d\lambda_\mu^a}{2\pi} \Big] \int_{-\infty}^{\infty} \Big[ \prod_a dy_\mu^a \Big] \exp\Big[ i \sum_a (\lambda_\mu^a - \vartheta) y_\mu^a + i d_1^{inp} \sum_a y_\mu^a m^a - \frac{d_3^{inp}}{2} \Big( \sum_a (y_\mu^a)^2 + 2 \sum_{a<b} y_\mu^a y_\mu^b q^{ab} \Big) \Big] \\
&\equiv I_2(q^{ab}, m^a).
\end{aligned} \tag{12}$$

Using Eq. (10) and (12), the quantity $M(q^{ab}, m^a)$ defined through Eq. (7) can be written as

$$M(q^{ab}, m^a) = \frac{p}{C} \log \left[ \langle (1 - \delta_{\eta^\mu,0}) I_1(q^{ab}, m^a, \eta^\mu) + \delta_{\eta^\mu,0} I_2(q^{ab}, m^a) \rangle_{\eta^\mu} \right]. \tag{13}$$

We now insert Eq. (13) back to Eq. (4) and enforce the definitions of $m$ and $q$ in Eq. (11) using the identities

$$\begin{aligned}
1 &= C \int \frac{dq^{ab} d\hat{q}^{ab}}{2i\pi} \exp\left( -C \hat{q}^{ab} q^{ab} + \hat{q}^{ab} \sum_j J_j^a J_j^b \right) \\
1 &= \sqrt{C} \int \frac{dm^a d\hat{m}^a}{2i\pi} \exp\left( -\sqrt{C} \hat{m}^a m^a + \hat{m}^a \sum_j J_j^a \right)
\end{aligned} \tag{14}$$

and the normalization of Eq. (4) using

$$\delta\left( \sum_j {J_j^a}^2 - C \right) = \int \frac{dE^a}{4i\pi} \exp\Big( -\frac{E^a}{2} \sum_{j \neq i} {J_j^a}^2 + \frac{C E^a}{2} \Big) \tag{15}$$

such that the numerator in Eq. (4) can be written as

$$A = \int \Big[\prod_a \frac{dE^a}{4i\pi}\Big]\Big[\prod_a \sqrt{C}\frac{dm^a d\hat{m}^a}{2i\pi}\Big]\Big[\prod_{a<b} C\frac{dq^{ab}d\hat{q}^{ab}}{2i\pi}\Big]$$
$$e^{C[M(q,m)-\frac{1}{\sqrt{C}}\sum_a \hat{m}^a m^a - \sum_{a<b}\hat{q}^{ab}q^{ab}+\sum_a \frac{E^a}{2}]}\int\Big[\prod_{j,a} dJ^a_{ij}\Big]e^{-\sum_{a,j}\frac{E^a}{2}(J^a_j)^2+\sum_{a,j}\hat{m}^a J^a_j+\sum_{a<b}\hat{q}^{ab}J^a_{ij}J^b_{ij}}. \tag{16}$$

Defining the function

$$W(\hat{q}^{ab},\hat{m}^a,E^a) = \log\int\Big[\prod_a dJ^a\Big]\exp\Big(-\frac{1}{2}\sum_a E^a(J^a)^2+\sum_a \hat{m}^a J^a+\sum_{a<b}\hat{q}^{ab}J^aJ^b\Big) \tag{17}$$

we can write

$$A = \int \Big[\prod_a \frac{dE^a}{4i\pi}\Big]\Big[\prod_a \sqrt{C}\frac{dm^a d\hat{m}^a}{2i\pi}\Big]\Big[\prod_{a<b} C\frac{dq^{ab}d\hat{q}^{ab}}{2i\pi}\Big]e^{C[M(q^{ab},m^a)+W(\hat{q}^{ab},\hat{m}^a,E^a)-\frac{1}{\sqrt{C}}\sum_a \hat{m}^a m^a-\sum_{a<b}\hat{q}^{ab}q^{ab}+\sum_a\frac{E^a}{2}]} \tag{18}$$

We can then compute $A$ in Eq. (18) using the saddle point approximation, by maximizing the argument of the exponential, that is maximising

$$G(q^{ab},\hat{q}^{ab},m^a,\hat{m}^a,E^a) \equiv M(q^{ab},m^a)+W(\hat{q}^{ab},\hat{m}^a,E^a)-\frac{1}{\sqrt{C}}\sum_a \hat{m}^a m^a-\sum_{a<b}\hat{q}^{ab}q^{ab}+\sum_a\frac{E^a}{2}. \tag{19}$$

In order to proceed to make this extremisation we assume a replica symmetric ansatz:

$$\begin{aligned} q^{ab} &= q \\ \hat{q}^{ab} &= \hat{q} \\ m^a &= m \\ \hat{m}^a &= \hat{m} \\ E_a &= E \end{aligned} \tag{20}$$

with these assumptions

$$G(q,\hat{q},m,\hat{m},E) = M(q,m)+W(\hat{q},\hat{m},E)+\frac{n}{2}(-\frac{2\hat{m}m}{\sqrt{C}}+\hat{q}q+E). \tag{21}$$

In the above Eq. (21), $W$ and $M$ are calculated using the limits for $n\to 0$ of the expressions in Eq. (13) and (17), as follows. For $W$, we use the Gaussian trick

$$e^{-x^2/2} = \int\frac{dt}{\sqrt{2\pi}}e^{-t^2/2+itx} = \int\frac{dt}{\sqrt{2\pi}}e^{-t^2/2\pm tx} \tag{22}$$

combined with the replica symmetric expression for $W$ to get

$$\begin{aligned} W(\hat{m},\hat{q},E) &= \log\int\Big[\prod_a dJ^a\Big]\exp\Big(-\frac{E}{2}\sum_a(J^a)^2+\hat{m}\sum_a J^a+\frac{\hat{q}}{2}\Big(\sum_a J^a\Big)^2-\frac{\hat{q}}{2}\sum_a(J^a)^2\Big) \\ &= \log\int\frac{dt}{\sqrt{2\pi}}e^{-t^2/2}\left[\int dJ\exp\left(-\frac{E+\hat{q}}{2}J^2+(\hat{m}+\sqrt{\hat{q}}t)J\right)\right]^n \end{aligned} \tag{23}$$

Using $a^n \approx 1+n\log a$ and $log(1+a)\approx a$, we have

$$W(\hat{m},\hat{q},E) = n\int\frac{dt}{\sqrt{2\pi}}e^{-t^2/2}\log\left[\int dJ\exp\left(-\frac{E+\hat{q}}{2}J^2+(\hat{m}+\sqrt{\hat{q}}t)J\right)\right] \tag{24}$$

In order to perform the Guassian integrals one can show that for general $a$, $b$ parameters:

$$\begin{aligned} \int dx e^{ax^2\pm bx} &= \sqrt{\frac{\pi}{a}}e^{\frac{b^2}{4a}} \\ \int\frac{dx}{\sqrt{2\pi}}e^{-x^2/2}(a+bx)^2 &= a^2+b^2 \end{aligned}$$

Therefore, integrating over $J$ in Eq. (24), leads to:

$$W(\hat{m},\hat{q},E) = n\Big(\int \frac{dt}{\sqrt{2\pi}} e^{-t^2/2} \log\sqrt{\frac{2\pi}{E+\hat{q}}} + \int \frac{dt}{\sqrt{2\pi}} e^{-t^2/2} \frac{(\hat{m}+\sqrt{\hat{q}}t)^2}{2(E+\hat{q})}\Big) \tag{25}$$

and over $t$, finally leads to:

$$W(\hat{m},\hat{q},E) = \frac{n}{2}\Big[\log(2\pi) - \log(E+\hat{q}) + \frac{\hat{q}+\hat{m}^2}{E+\hat{q}}\Big] \tag{26}$$

Computing $M$ is a bit more tricky.

$$M(q,m) = \frac{p}{C}\log\left[\langle(1-\delta_{\eta^\mu,0})I_1(q,m,\eta^\mu) + \delta_{\eta^\mu,0}I_2(q,m)\rangle_{\eta^\mu}\right]. \tag{27}$$

as one have to compute $I_1(q,m,\eta^\mu)$ and $I_2(q,m)$. Using the Gaussian trick in Eq. (22) and assuming replica symmetry we rewrite Eq. (10) as

$$\begin{aligned} I_1(q,m,\xi) &= \int_{-\infty}^{\infty}\Big[\prod_a \frac{dx_\mu^a}{2\pi}\Big]\exp\Big\{\Big[-\frac{i}{g}(\eta^\mu+g\vartheta)+id_1^{inp}m\Big]\sum_a x_\mu^a - \frac{d_3^{inp}}{2}\sum_a (x_\mu^a)^2 + -d_3^{inp}q\sum_{a<b}x_\mu^a x_\mu^b\Big\} \\ &= \int_{-\infty}^{\infty}\Big[\prod_a \frac{dx_\mu^a}{2\pi}\Big]\exp\Big\{\Big[-\frac{i}{g}(\eta^\mu+g\vartheta)+id_1^{inp}m\Big]\sum_a x_\mu^a - \frac{d_3^{inp}}{2}\sum_a (x_\mu^a)^2 + \frac{d_3^{inp}q}{2}\sum_a (x_\mu^a)^2 - \frac{qd_3^{inp}}{2}\Big(\sum_a x_\mu^a\Big)^2\Big\} \\ &= \int Dt\Big\{\int \frac{dx_\mu}{2\pi}\exp\Big[-i\left(g^{-1}\eta^\mu+\vartheta-d_1^{inp}m-t\sqrt{qd_3^{inp}}\right)x_\mu - \frac{d_3^{inp}}{2}(1-q)x_\mu^2\Big]\Big\}^n \end{aligned} \tag{28}$$

with $Dt = \frac{dt}{\sqrt{2\pi}}e^{-t^2/2}$. In a very similar way we can write Eq. (12) as

$$I_2(q,m) = \int Dt\Big\{\int_0^\infty \frac{d\lambda_\mu}{2\pi}\int_{-\infty}^{\infty} dy_\mu \exp\Big[i\left(\lambda_\mu^a - \vartheta + d_1^{inp}m + t\sqrt{qd_3^{inp}}\right)y_\mu - \frac{d_3^{inp}}{2}(1-q)(y_\mu)^2\Big]\Big\}^n. \tag{29}$$

We define $P(\eta^\mu>0)=f$ and rewrite Eq. (13) as

$$\begin{aligned} M(q,m) &= \frac{p}{C}\log\{\langle(1-\delta_{\eta^\mu,0})\rangle_{\eta^\mu}\langle I_1(q,m,\eta^\mu)\rangle_{\eta^\mu} + \langle\delta_{\eta^\mu,0}\rangle_{\eta^\mu}I_2(q,m)\} \\ &= \frac{p}{C}\log\Big[f\langle I_1(q,m,\eta^\mu)\rangle_{\eta^\mu} + (1-f)I_2(q,m)\Big]. \end{aligned} \tag{30}$$

Simplifying for the sake of visualization Eq. (28) and (29) as

$$\begin{aligned} I_1(q,m,\eta^\mu) &= \int DtY^n \\ I_2(q,m) &= \int DtK^n \end{aligned} \tag{31}$$

where

$$\begin{aligned} Y &\equiv \int \frac{dx_\mu}{2\pi}\exp\Big[-i\left(g^{-1}\eta^\mu+\vartheta-d_1^{inp}m-t\sqrt{qd_3^{inp}}\right)x_\mu - \frac{d_3^{inp}}{2}(1-q)x_\mu^2\Big] \\ K &\equiv \int_0^\infty \frac{d\lambda_\mu}{2\pi}\int_{-\infty}^{\infty} dy_\mu\exp\Big[i\left(\lambda_\mu^a-\vartheta+d_1^{inp}m+t\sqrt{qd_3^{inp}}\right)y_\mu - \frac{d_3^{inp}}{2}(1-q)(y_\mu)^2\Big] \end{aligned} \tag{32}$$

one can use again $a^n \approx 1 + n\log a$ and $\log(1+a)\approx a$, which is valid for $n\to 0$, to write $M(q,m)$ as

$$\begin{aligned} M(q,m) &= \frac{p}{C}\log\Big[f\Big\langle\int DtY^n\Big\rangle_{\eta^\mu} + (1-f)\int DtK^n\Big] \\ &= \frac{p}{C}\log\Big[\int Dt[f\langle 1+n\log Y\rangle_{\eta^\mu} + (1-f)(1+n\log K)\Big] \\ &= \frac{p}{C}\log\Big[1+n\Big(f\int Dt\langle\log Y\rangle_{\eta^\mu} + (1-f)\int Dt\log K\Big)\Big] \\ &= \frac{p}{C}n\Big(f\int Dt\langle\log Y\rangle_{\eta^\mu} + (1-f)\int Dt\log K\Big) \end{aligned} \tag{33}$$

Turning back to the original notation we can further develop the terms composing the above approximation. The first one yields:

$$\begin{aligned}
\int Dt\,\langle\log Y\rangle_{\eta^\mu} &= \int Dt \int \left\langle \frac{dx_\mu}{2\pi}\exp\Big[-i\left(g^{-1}\eta^\mu+\vartheta-d_1^{inp}m-t\sqrt{qd_3^{inp}}\right)x_\mu-\frac{d_3^{inp}}{2}(1-q)x_\mu^2\Big]\right\rangle_{\eta^\mu}\\
&= \int Dt \left\langle \log\Big[\exp\Big\{-\frac{\left(d_1^{inp}m-g^{-1}\eta^\mu-\vartheta+t\sqrt{qd_3^{inp}}\right)^2}{2d_3^{inp}(1-q)}\Big\}\sqrt{\frac{2\pi}{d_3^{inp}(1-q)}}\frac{1}{2\pi}\Big]\right\rangle_{\eta^\mu} \qquad (34)\\
&= \frac{1}{2}\left[-\log 2\pi-\log d_3^{inp}(1-q)-\frac{\left\langle\left(d_1^{inp}m-g^{-1}\eta^\mu-\vartheta\right)^2\right\rangle_{\eta^\mu}+qd_3^{inp}}{d_3^{inp}(1-q)}\right]
\end{aligned}$$

and the second one yields:

$$\begin{aligned}
\int Dt\log K &= \int Dt\log\int_0^\infty\frac{d\lambda_\mu}{2\pi}\int_{-\infty}^\infty dy_\mu\exp\left[i\left(\lambda_\mu^a-\vartheta+d_1^{inp}m+t\sqrt{qd_3^{inp}}\right)y_\mu-\frac{d_3^{inp}}{2}(1-q)(y_\mu)^2\right]\\
&= \int Dt\log\int_0^\infty\frac{d\lambda_\mu}{2\pi}\exp\left[-\frac{\left(d_1^{inp}m+\lambda_{\mu-\vartheta+t\sqrt{qd_3^{inp}}}\right)^2}{1d_3^{inp}(1-q)}\right]\sqrt{\frac{2\pi}{d_3^{inp}(1-q)}} \qquad (35)\\
&= \int Dt\log\int_{\frac{d_1^{inp}m-\vartheta+t\sqrt{qd_3^{inp}}}{\sqrt{d_3^{inp}(1-q)}}}^\infty\frac{dz}{\sqrt{2\pi}}e^{\frac{-z^2}{2}}
\end{aligned}$$

where in the last passage we made a simple change of variables. Therefore we can rewrite Eq. (30) as:

$$\begin{aligned}
M(q,m) &= \frac{p}{C}n\left\{\frac{f}{2}\left[-\log[2\pi d_3^{inp}(1-q)]-\frac{\left[\left\langle\left(d_1^{inp}m-g^{-1}\eta^\mu-\vartheta\right)^2\right\rangle_{\eta^\mu}+qd_3^{inp}\right]}{d_3^{inp}(1-q)}\right]+(1-f)\int Dt\log H(u)\right\}\\
&\text{where}\\
u &\equiv \frac{d_1^{inp}m-\vartheta+t\sqrt{qd_3^{inp}}}{\sqrt{d_3^{inp}(1-q)}}\\
H(u) &\equiv \int_u^\infty\frac{dt}{\sqrt{2\pi}}e^{-t^2/2}.
\end{aligned} \qquad (36)$$

Now we can evaluate the derivatives

$$\frac{dG}{d\hat{m}}=\frac{dG}{d\hat{q}}=\frac{dG}{dE}=\frac{dG}{dm}=\frac{dG}{dq}=0 \qquad (37)$$

where $G=G(q,\hat{q},m,\hat{m},E)$ given by Eq. (21), and set them to zero to find the maximum of Eq. (21), with $W(\hat{m},\hat{q},E)$ given by Eq. (26) and $M(q,m)$ given by Eq. (36).
With the first three derivatives equalized to zero, which are applied only to the second and third term of Eq. (21), and assuming $Cq\gg m^2$ and $|C(1-2q)|\gg m^2$ as $C\to\infty$, we obtain the relations

$$\begin{aligned}
\hat{m} &= -\frac{m}{\sqrt{C}(q-1)}\\
\hat{q} &= \frac{q}{(1-q)^2} \qquad (38)\\
E &= \frac{1-2q}{(q-1)^2}.
\end{aligned}$$

Substituting them into Eq. (21) we have to perform the last two derivatives.
$\frac{dG}{dm}$ can be simply evaluated, applying the Leibniz integral rule $\frac{d}{dx}H(f(x)) = \frac{d}{dx}\int_{f(x)}^{\infty}\frac{dt}{\sqrt{2\pi}}e^{\frac{-t^2}{2}} = -\exp(-\frac{f(x)^2}{2})\frac{d}{dx}f(x)$ yielding:

$$\frac{dG}{dm} = 0 = -fd_1^{inp}(d_1^{inp}m - g^{-1}\langle\eta^\mu\rangle - \vartheta) - \frac{\sqrt{d_3^{inp}(1-q)}(1-f)d_1^{inp}}{\sqrt{2\pi}}\int DtH(u)^{-1}e^{-u^2/2} \tag{39}$$

The derivative in $q$ requires in addition the integration by parts of the term multiplied by $(1-f)$ enabling to reach the simplified solution:

$$\frac{dG}{dq} = 0 = \frac{\alpha}{q}\left\{f\left[\frac{\langle(d_1^{inp}m - g^{-1}\eta^\mu - \vartheta)^2\rangle + qd_3^{inp}}{d_3^{inp}}\right] + \frac{(1-f)(1-q)}{2\pi}\int DtH(u)^{-2}e^{-u^2}\right\} \tag{40}$$

where $\alpha \equiv p/C$ is the storage capacity.
As explained in the main text we take the limit $q \to 1$, in which the storage capacity $\alpha$ becomes the critical one $\alpha_c$. Note that in this limit:

$$\lim_{q\to 1} u = \begin{cases} \infty & \text{if } t > \frac{\vartheta - d_1^{inp}m}{\sqrt{d_3^{inp}}} \\ -\infty & \text{if } t < \frac{\vartheta - d_1^{inp}m}{\sqrt{d_3^{inp}}}. \end{cases} \tag{41}$$

This enables to further simplify the above equations as

$$\lim_{x\to-\infty} H(x) \approx 1$$
$$\lim_{x\to\infty} H(x) \approx \frac{1}{\sqrt{2\pi}u}e^{-u^2/2}(1 - \frac{1}{u^2}) = \frac{1}{\sqrt{2\pi}u}e^{-u^2/2}$$

where in the second approximation we have Taylor expanded $H(x)$ around $x = 0$.
The simple application of the limit $q \to 1$ with the above approximations, and the introduction of the variable $x = \frac{\vartheta - d_1^{inp}m}{\sqrt{d_3^{inp}}}$ leads to the final set of equations for the critical storage capacity

$$\begin{cases} f(x + \frac{d_1^{out}}{g\sqrt{d_3^{inp}}}) = (1-f)\int_x^\infty Dt(t-x) \\ \frac{1}{\alpha_c} = f\left[x^2 + \frac{d_2^{out}}{g^2 d_3^{inp}} + \frac{2xd_1^{out}}{g\sqrt{d_3^{inp}}} + 1\right] + (1-f)\int_x^\infty Dt(t-x)^2. \end{cases} \tag{42}$$

where $d_{1,2,3}^{out}$ are defined in the same way as $d_{1,2,3}^{inp}$ except that the averages are now over the output distribution $\eta$.

Going from the calculation reported above for the threshold-linear perceptron it is straightforward to calculate the optimal capacity of a network of threshold linear units. Considering the network defined through Eq. (1) of the main text, the corresponding volume we need to calculate can be written as

$$V_T = \frac{\int\prod_{i,j,j\neq i} dJ_{ij}\delta\left(\sum_{j,j\neq i}J_{ij}^2 - C\right)\prod_{i,\mu}\left[(1-\delta_{\eta^\mu,0})\,\delta\left(h_i^\mu - \vartheta - \frac{\eta^\mu}{g}\right) + \delta_{\eta^\mu,0}\Theta\left(\vartheta - h_i^\mu\right)\right]}{\int\prod_{i,j,j\neq i} dJ_{ij}\prod_i\delta\left(\sum_{j,j\neq i}J_{ij}^2 - C\right)} \tag{43}$$

Since $V_T$ can be written as the product of the individual volumes of the connection weights towards each unit, as $V_T = \prod_i^N V_i$ and thus $\langle \log V_T\rangle_\eta = N\langle \log V_i\rangle_\eta$, we will essentially be dealing with individual perceptrons like the one we just studied. Putting $d_1^{inp} = d_1^{out} = d_1$ and $d_2^{inp} = d_2^{out} = d_2$ and thus $d_3^{inp} = d_3^{out} = d_3$ for $\forall i$, we arrive to the equations presented in Eq. (3) of the main text.
As explained in the main text, we evaluate the maximal storage capacity in the limit $g \to \infty$, which is reached for moderate values of $g$. Eq. (3) of the main text in the $g \to \infty$ limit reduces to:

$$\begin{cases} 0 = fx - (1-f)\int_x^\infty Dt(t-x) \\ \frac{1}{\alpha_c} = f(x^2+1) + (1-f)\int_x^\infty Dt(t-x)^2, \end{cases} \tag{44}$$

which provides the universal $\alpha_c^G$ bound for errorless retrieval, dependent only through $f$ on the distribution of the patterns.

#### B. Derivation of the limits

From Eq. (3) of the main text it is possible to evaluate the two limits of very sparse and non-sparse coding. First, a simple substitution at $f = 1$ leads to

$$x = -\frac{d_1}{g\sqrt{d_3}} \tag{45}$$

$$\alpha_c^{-1} = 1 + \frac{1}{g^2}. \tag{46}$$

The case $f \to 0$ is a bit trickier. We first rearrange the first equation in Eq. (3) as

$$\frac{f}{1-f} = \frac{1}{(x + \frac{d_1}{g\sqrt{d_3}})}\int_x^\infty Dt(t-x) = \frac{1}{(x + \frac{d_1}{g\sqrt{d_3}})}\Big(\frac{e^{-\frac{x^2}{2}}}{\sqrt{2\pi}} - x\int_x^\infty Dt\Big) \tag{47}$$

As $f$ goes to zero, for the left hand side to be equal to the right hand side, we should have $x \to \infty$. We therefore use the expansion

$$\int_x^\infty Dt = \frac{e^{-\frac{x^2}{2}}}{\sqrt{2\pi}}\Big[\frac{1}{x} - \frac{1}{x^3} + \mathcal{O}\Big(\frac{1}{x^5}\Big)\Big]$$

to write the right hand side of Eq. (47) as

$$\frac{f}{1-f} \approx \frac{e^{-\frac{x^2}{2}}}{\sqrt{2\pi}x^3}. \tag{48}$$

We find a solution to Eq. (48) through the following iterative procedure. We first solve the leading term for $f \to 0$ in $x \to \infty$ namely

$$f \approx \frac{e^{-\frac{x^2}{2}}}{\sqrt{2\pi}}.$$

yielding

$$x \approx \sqrt{2\ln\Big(\frac{1}{\sqrt{2\pi}f}\Big)} \tag{49}$$

We then insert $x$ from Eq. (49) into $\exp(-x^2/2) = \sqrt{2\pi}fx^3$ to obtain the logarithmic correction

$$\begin{aligned} e^{-\frac{x^2}{2}} &\approx \sqrt{2\pi}fx^3 \\ x &\approx \sqrt{2\ln\Big(\frac{1}{\sqrt{2\pi}fx^3}\Big)} \\ x &\approx \sqrt{2\ln\Big(\frac{1}{\sqrt{2\pi}f}\Big)\Big(1 - \frac{\ln x^3}{\ln\frac{1}{\sqrt{2\pi}f}}\Big)} \\ &\approx \sqrt{2\ln\Big(\frac{1}{\sqrt{2\pi}f}\Big)}\left(1 - \frac{3}{4}\frac{\ln\Big(2\ln(\frac{1}{\sqrt{2\pi}f})\Big)}{\ln\frac{1}{\sqrt{2\pi}f}}\right). \end{aligned} \tag{50}$$

where in the last passage we have used the Taylor expansion of the square $\sqrt{1-y} = 1 - \frac{y}{2} + \mathcal{O}(y^2)$ around $y = 0$ as for $f \to 0$, $\frac{\ln x^3}{\ln\frac{1}{\sqrt{2\pi}f}} \to 0$.

We have tested numerically that the above expression Eq. (50) for $x$ is indeed a solution to Eq. (47) for $f \to 0$.

We now proceed to evaluating $\alpha_c$, we apply the same Taylor expansion as before

$$\begin{aligned}
\alpha_c &= \Big\{ f[\langle (x+\frac{\xi_i}{g\sqrt{d_3}})\rangle^2 + 1] + (1-f)\int_x^\infty Dt(t-x)^2 \Big\}^{-1} \\
&= \Big\{ f[\langle (x+\frac{\xi_i}{g\sqrt{d_3}})\rangle^2 + 1] + (1-f)\Big( -\frac{xe^{-\frac{x^2}{2}}}{\sqrt{2\pi}} + (1+x^2)\int_x^\infty Dt \Big) \Big\}^{-1} \\
&\approx \Big\{ fx^2 - \frac{xe^{-\frac{x^2}{2}}}{\sqrt{2\pi}} + \frac{(1+x^2)}{\sqrt{2\pi}} e^{-\frac{x^2}{2}} \Big( \frac{1}{x} - \frac{1}{x^3} + \frac{3}{x^5} \Big) \Big\}^{-1} \\
&\approx \Big\{ fx^2 + \frac{e^{-\frac{x^2}{2}}}{\sqrt{2\pi}} \Big( -x + \frac{(1+x^2)(x^4 - x^2 + 3)}{x^5} \Big) \Big\}^{-1} \\
&\approx \Big\{ fx^2 + \frac{e^{-\frac{x^2}{2}}}{\sqrt{2\pi}} \Big( \frac{2x^2+3}{x^5} \Big) \Big\}^{-1} = \Big\{ fx^2 + \sqrt{\frac{2}{\pi}} \frac{e^{-\frac{x^2}{2}}}{x^3} \Big\}^{-1}.
\end{aligned}$$

To summarise in the limit $f \to 0$ we obtain

$$\begin{cases}
x \approx \sqrt{2\ln\Big(\frac{1}{\sqrt{2\pi}f}\Big)} \left( 1 - \frac{3}{4} \frac{\ln\Big(2\ln(\frac{1}{\sqrt{2\pi}f})\Big)}{\ln \frac{1}{\sqrt{2\pi}f}} \right) \\
\alpha_c \approx \Big\{ fx^2 + \sqrt{\frac{2}{\pi}} \frac{e^{-\frac{x^2}{2}}}{x^3} \Big\}^{-1}.
\end{cases} \tag{51}$$

Substituting $x$ in $\alpha_c$ to the leading order leads to Eq.(5) presented in the main text.

### C. Training algorithm

For the purpose of assessing whether the Gardner capacity for errorless retrieval can be reached with explicit training, we can decompose a network of, say, $N+1 = 10001$ units into $N+1$ independent threshold linear perceptrons. A threshold linear perceptron is just a 1-layer feedforward neural network with $N$ inputs and one output, the activity of which is given by a threshold-linear *activation function.*

$$[h]^+ = \max(0, h) \tag{52}$$

The network is trained with $p$ patterns. One can then think of the input as a matrix $\bar{\xi}$ of dimension $[N \times p]$ and of the output as a vector $\vec{\eta}$ of dimension $[1 \times p]$.

The aim of the algorithm is to tune the weights such that all $p$ patterns can be memorized. In order to tune the weights we start from an initial connectivity vector $\vec{J_0}$ of dimension $[1 \times N]$ and estimate the output $\vec{\hat{\eta}}$ as:

$$\begin{aligned}
\vec{h} &= \vec{J}\bar{\xi} \\
\vec{\hat{\eta}} &= g[\vec{h}]^+
\end{aligned} \tag{53}$$

where $g$ is the gain parameter. We then compare the output $\vec{\hat{\eta}}$ with the desired output $\vec{\eta}$ through the loss function

$$\mathrm{L}(\vec{\hat{\eta}}) = \sum_{\mu=1}^{p} \frac{1}{2} (\hat{\eta}^\mu - \eta^\mu)^2. \tag{54}$$

The TL perceptron algorithm can be seen as simply a stripped down version of *backpropagation*, for a 1-layer network: the weights $\vec{J}$ are modified by gradient descent to minimize the loss during the steps $k = 1..k^{MAX}$ where $k^{MAX}$ is the number of steps needed for the gradient descent in order to reach the minima $\frac{d\mathrm{L}(\vec{J_k})}{d\vec{J_k}} = 0$. If at the minima $\mathrm{L}(\tilde{\mathrm{J}}_{\mathrm{k}^{\mathrm{MAX}}}) = 0$ at least a set of weights exists for errorless retrieval at that $p$ value. The storage capacity $\alpha_c = \frac{p^{max}}{N}$ is evaluated by estimating $p^{max}$ as the highest $p$ value enabling to reach $\mathrm{L}(\tilde{\mathrm{J}}_{\mathrm{k}^{\mathrm{MAX}}}) = 0$.

Initializing the weights around zero facilitates reaching the minima. The chain derivative that in general implements gradient descent in backpropagation, in this case reduces to

$$\vec{J}_{k+1} = \vec{J_k} + \gamma \frac{g}{p} (\vec{\eta} - \vec{\hat{\eta}}) \Theta(\vec{\hat{\eta}}) \bar{\xi}^T \tag{55}$$

where $\Theta(\vec{\hat{\eta}})$ is the Heaviside step function applied to all $N$ elements of $\vec{\hat{\eta}}$ and where $\gamma$ is a learning rate, which we vary in order to facilitate reaching the minima.

#### D. Hebbian capacity and sparsity of the retrieved pattern

From the calculation reported in [1], it can be shown that for a network of threshold-linear units described in Eq. (1) of the main text in which $p$ patterns are stored through Hebbian learning, the storage capacity $\alpha_c$ can be found as the value of $\alpha$ for which there are values of $v_c$ and $w_c$ that solve the equation

$$A_2(w,v)^2 - \alpha A_3(w,v) = 0 \tag{56}$$

at a single point on the $w, v$ plane; where

$$A_2(w,v) = \frac{a}{v(1-a)}\Big\langle\Big(\frac{\eta}{\langle\eta\rangle} - 1\Big)(x\phi(x) + \sigma(x)\Big\rangle \tag{57}$$

$$A_3(w,v) = \Big\langle(x^2+1)\phi(x) + \sigma(x)\Big\rangle \tag{58}$$

and

$$x \equiv w + v\frac{\eta}{\langle\eta\rangle} \tag{59}$$

$$\phi(x) = \frac{[1+\mathrm{erf}(\frac{x}{\sqrt{2}})]}{2} = \frac{\mathrm{erfc}(\frac{-x}{\sqrt{2}})}{2} \tag{60}$$

$$\sigma(x) = \frac{e^{-x^2/2}}{\sqrt{2\pi}}. \tag{61}$$

and the auxiliary variables $v$ and $w$, defined as in [2] and dependent on the threshold $\vartheta$, quantify the signal to noise ratio respectively of the specific signal of the pattern to be retrieved and the one of the background both versus the noise due to memory loading [1]. Estimating $v_c$ and $w_c$ translates to optimizing the threshold $\vartheta$ such that it maximizes the storage capacity.

In [1] the above expression are reported assuming $a = \langle\eta\rangle = \langle\eta^2\rangle$, but the above equations do not make this assumption and can be derived easily from the calculation reported in [1].

Following [1], the average of the activity and the average of the square activity in the patterns retrieved with Hebbian weights are calculated considering that the field, i.e. the input received by a cell with activity $\eta$ in the memory, is normally distributed around a mean field proportional to $x$. If we call $z$ a random variable normally distributed with mean zero and variance one, $x$ is already the mean field properly normalized. With the threshold-linear transfer function, the output will be $g(x+z)$ for $x+z>0$ and 0 with probability $\phi(-x)$. Therefore the average activity and the average square activity are:

$$\langle V\rangle = g\langle\int_{-x_c(\eta)}^{\infty} Dz[x_c(\eta)+z]\rangle_\eta = g\langle[x_c\phi(x_c)+\sigma(x_c)]\rangle_\eta \tag{62}$$

$$\langle V^2\rangle = g^2\langle\int_{-x_c(\eta)}^{\infty} Dz[x_c(\eta)+z]^2\rangle_\eta = g^2\langle[(1+x_c^2)\phi(x_c)+x_c\sigma(x_c)]\rangle_\eta \tag{63}$$

$$x_c \equiv w_c + v_c\frac{\eta}{\langle\eta\rangle}, \tag{64}$$

the sparsity of the retrieved memory is thus $a_r^H = \langle V\rangle^2/\langle V^2\rangle$.

As reported in the main text, we have compared capacity values using a binary, ternary, quaternary and an exponential distribution:

$$p(\eta) = (1-a)\delta(\eta) + a\delta(1-x) \tag{65}$$

$$p(\eta) = (1-\frac{9a}{5})\delta(\eta) + \frac{3a}{2}(\eta-\frac{1}{3}) + \frac{3a}{10}\delta(\eta-\frac{5}{3}) \tag{66}$$

$$p(\eta) = (1-\frac{9a}{4})\delta(\eta) + \frac{3a}{2}\delta(\eta-\frac{2}{9}) + \frac{3a}{5}\delta(\eta-\frac{5}{9}+\frac{3a}{20}(\eta-\frac{20}{9}) \tag{67}$$

$$P(\eta) = (1-2a)\delta(\eta) + 4a\exp(-2\eta) \tag{68}$$

One can see that all distributions are such that $\langle\eta\rangle = \int_0^\infty d\eta P(\eta)\eta = a$ and $\langle\eta^2\rangle = \int_0^\infty d\eta P(\eta)\eta^2 = a$, so that $a$ coincides with the sparsity $\langle\eta\rangle^2/\langle\eta^2\rangle$ of the network. The fraction of active units is thus related to $a$ as $f = a, 9a/5, 9a/4$, $2a$ respectively.

As a supplement to Fig. 2 of the main text, reproduced here in the 3 separate panels in the upper row in Fig. 1, we show a comparison between the Hebbian capacity and the Gardner one when plotted as a function of the output sparsity (in the bottom row of Fig. 1). The Gardner storage capacity is now in each of these 3 cases above the Hebbian capacity, taken as a function of the output sparsity instead of the input one.

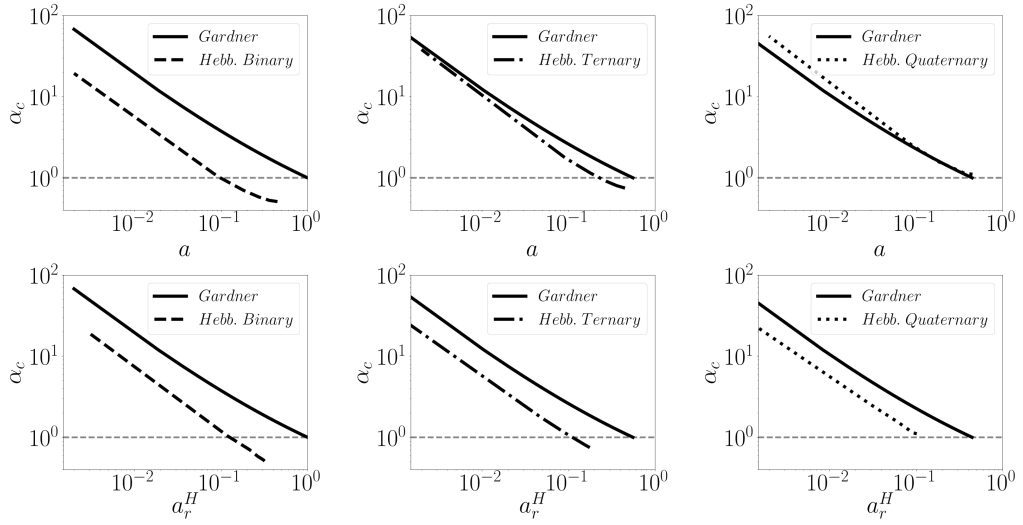


FIG. 1. Suplementary to Fig. (2). Comparison between the Hebbian and Gardner storage capacity for 3 discrete distributions. The upper row considers as sparsity parameter the one of the input pattern, the lower row the one of the retrieved pattern. The Garner capacity is that given by Eq. (3) of the main text

### E. Analytical derivation for the exponential distribution

In order to facilitate the comparison we extend the analytical calculations in [1] and evaluate explicitly the analytical expression of Eq. (57) and (58) for the exponential distribution. In general, for $A_2$ we write

$$\begin{aligned} A_2 &= \frac{a}{v(1-a)} \int_0^\infty d\eta P(\eta)(\frac{\eta}{\langle\eta\rangle} - 1) \int_{-\infty}^{x(\eta)} Dz(x(\eta) - z) \\ &= \frac{a}{v(1-a)} \Big\{ \int_w^\infty Dz \int_{\frac{(z-w)\langle\eta\rangle}{v}}^\infty d\eta P(\eta)(\frac{\eta}{\langle\eta\rangle} - 1)(x(\eta) - z) + \int_{-\infty}^w Dz \int_0^\infty d\eta P(\eta)(\frac{\eta}{\langle\eta\rangle} - 1)(x(\eta) - z) \Big\} \end{aligned} \tag{69}$$

with $x(\eta) \equiv w + v\eta/\langle\eta\rangle$. Substituting Eq. (68) we obtain

$$\begin{aligned} A_2^{exp} &= \frac{a}{v(1-a)}(A_{2.1} + A_{2.2} + A_{2.3}) \\ A_{2.1} &= \int_{-\infty}^w Dz \int_0^\infty d\eta 4a \exp(-2\eta)(\frac{\eta}{a} - 1)(w + \frac{v\eta}{a} - z) \\ A_{2.2} &= \int_{-\infty}^w Dz(1-2a)(z-w) \\ A_{2.3} &= \int_w^\infty Dz \int_{\frac{(z-w)a}{v}}^\infty d\eta 4a \exp(-2\eta)(\frac{\eta}{a} - 1)(w + \frac{v\eta}{a} - z). \end{aligned} \tag{70}$$

Solving the equations leads to

$$\begin{aligned} A_{2.1} &= (1-2a)\sigma(w) + \Big[\frac{v}{a} + w - v - 2wa\Big]\phi(w) \\ A_{2.2} &= (2a-1)(\sigma(w) + w\phi(w)) \\ A_{2.3} &= \exp\Big(\frac{2aw}{v}\Big)\exp\Big(\frac{2a^2}{v^2}\Big)\Big[\frac{v(1-a)}{a}\phi\Big(-w-\frac{2a}{v}\Big) + \sigma\Big(w+\frac{2a}{v}\Big) - \Big(w+\frac{2a}{v}\Big)\phi\Big(-w-\frac{2a}{v}\Big)\Big]. \end{aligned} \tag{71}$$

Thus

$$A_2^{exp} = \phi(w) + \exp\Big(\frac{2aw}{v} + \frac{2a^2}{v^2}\Big)\Big\{\phi\Big(-w-\frac{2a}{v}\Big) + \frac{a}{v(1-a)}\Big[\sigma\Big(w+\frac{2a}{v}\Big) - \Big(w+\frac{2a}{v}\Big)\phi\Big(-w-\frac{2a}{v}\Big)\Big]\Big\} \tag{72}$$

For $A_3$ we have

$$\begin{aligned} A_3^{exp} &= A_{3.1} + A_{3.2} + A_{3.3} \\ A_{3.1} &= \int_{-\infty}^{w} Dz \int_0^{\infty} d\eta 4a \exp(-2\eta)(w + \frac{v\eta}{a} - z)^2 \\ A_{3.2} &= \int_{-\infty}^{w} Dz(1-2a)(w-z)^2 \\ A_{3.3} &= \int_{w}^{\infty} Dz \int_{\frac{(z-w)a}{v}}^{\infty} d\eta 4a \exp(-2\eta)(w + \frac{v\eta}{a} - z)^2 \end{aligned} \tag{73}$$

Substituting Eq. (68) we obtain

$$\begin{aligned} A_{3.1} &= (1-2a)\sigma(w) + \Big[\frac{v}{a} + w - v - 2wa\Big]\phi(w) \\ A_{3.2} &= (2a-1)(\sigma(w) + w\phi(w)) \\ A_{3.3} &= \exp\Big(\frac{2aw}{v}\Big)\exp\Big(\frac{2a^2}{v^2}\Big)\Big[\frac{v(1-a)}{a}\phi\Big(-w-\frac{2a}{v}\Big) + \sigma\Big(w+\frac{2a}{v}\Big) - \Big(w+\frac{2a}{v}\Big)\phi\Big(-w-\frac{2a}{v}\Big)\Big] \end{aligned} \tag{74}$$

and solving the equations leads to

$$\begin{aligned} A_{3.1} &= 2a\Big[\sigma(w)(w+\frac{v}{a}) + \phi(w)(1 + w^2 + \frac{vw}{a} + \frac{v^2}{2a^2}\Big] \\ A_{3.2} &= (1-2a)[w\sigma(w) + (1+w^2)\phi(w))] \\ A_{3.3} &= \frac{v^2}{a}\exp\Big(\frac{2aw}{v}\Big)\exp\Big(\frac{2a^2}{v^2}\Big)\phi\Big(-w-\frac{2a}{v}\Big). \end{aligned} \tag{75}$$

Thus

$$A_3^{exp} = 2v(\sigma(w) + \phi(w)) + w\sigma(w) + (1+w^2)\phi(w) + \frac{v^2}{a}\phi(w) + \exp\Big(\frac{2aw}{v} + \frac{2a^2}{v^2}\Big)\phi(-w - \frac{2a}{v}). \tag{76}$$

#### F. Comparison with real data

In the real activity distributions we use, each neuron emits, in time bins of fixed duration (we use 100msec), $0, \ldots, n, \ldots, n_{max}$ spikes, with relative frequency $c_n$, such that $\sum_{n=0}^{n_{max}} c_n = 1$. These values are taken from Fig. 2 of [3] and correspond to the histograms in blue in Fig.2 below (and in Fig.3 of the main text); they are assumed to be the distributions of the patterns to be stored. If the weights are those described by the Gardner calculation, these patterns can be retrieved as they are, and their distribution remains the same. If they are stored with Hebbian weights close to the maximal Hebbian capacity, however, the retrieved distributions look different, and they can be derived as follows.

The firing rate $V$ of a neuron in retrieving a stored pattern $\eta$ is assumed proportional to $w + v\eta/\langle\eta\rangle + z$ [1], where the parameters $w$ and $v$ are appropriately rescaled signal-to-noise ratios (general and pattern-specific), such that the normally distributed random variable $z$, of zero mean and unitary variance, is taken to describe all other non constant (noise) terms, besides $\eta$ itself. Averaging over $z$ one can write, as in Eq.(62), that at the maximal capacity

$$\langle V\rangle(\eta) = g\int_{-x_c(\eta)}^{\infty} Dz[x_c(\eta) + z] = g[x_c\phi(x_c) + \sigma(x_c)] \tag{77}$$

where $x \equiv w + v\eta/\langle\eta\rangle$ and at the saddle-point the parameters $w$ and $v$ take the values $w_c$ and $v_c$ that maximize capacity, as explained in [1]. This implies setting an optimal value for the threshold $\vartheta$, which in the analysis is absorbed into the parameter $w$, and which determines the sparsity of the retrieved distribution. The gain $g$ remains, however, a free parameter, that affects neither sparsity nor capacity. It is a rescaled version of the original gain $g$ in the hypothetical TL transfer function. In other words, the maximal Hebbian capacity determines the shape of the retrieval activity distribution, but not its scale (e.g., in spikes per sec).

To produce a histogram, that details the frequency with which the neuron would produce $n$ spikes at retrieval, e.g. again in bins of 100msec, one has to set this undetermined scale. We set it arbitrarily, with the rough requirement that the frequency of producing $n_{max}$ spikes at retrieval be below what it is in the observed distribution, taken to describe storage, and negligible for $n_{max}+1$ spikes. Having set the scale $g$, the frequency with which the neuron emits $n$ spikes at retrieval, with $0 < n < n_{max}$ is the probability that $n - 1/2 < V < n + 1/2$, that is, it is a sum over contributions from each $\eta$, such that

$$\begin{aligned} n - \frac{1}{2} < g(w_c + v_c \frac{\eta}{\langle\eta\rangle} + z) < n + \frac{1}{2} \\ \frac{n}{g} - \frac{1}{2g} - x_c < z < \frac{n}{g} + \frac{1}{2g} - x_c \end{aligned} \tag{78}$$

i.e.,

$$Pr(n) = \sum_{\eta=0}^{\eta_{max}} c_\eta \Big[\phi\Big(\frac{n}{g} + \frac{1}{2g} - x_c\Big) - \phi\Big(\frac{n}{g} - \frac{1}{2g} - x_c\Big)\Big], \tag{79}$$

with appropriate expressions for the two extreme bins. These are the distributions shown in Fig.3 in the main text, and in Fig.2 below.

We took $g = \frac{1}{2}$, as this value satisfies the a priori requirements and allows to keep the same number of bins in the retrieved memory as in the stored one (and the coefficients sum up to one, to a very good approximation).

#### *Analysis of the other recorded cells*

Supplementary to Fig. (3) in the main text, we report in Fig. 2 the same analysis for all 9 single cells reported (using 100ms bins) in [3].

In each panel we write the capacity *à la Gardner* and the Hebbian one (calculated without fitting an exponential) for the 9 empirical distributions, as well as the sparsity of the original distribution and the sparsity of the one that would be retrieved with Hebbian weights. For simplicity of visualization we also show the storage capacity values against each other, calculated *à la Gardner* and *à la Hebb* (again, without fitting an exponential), as a single scatterplot for the 9 distributions, in Fig. 3.

[1] A. Treves and E. T. Rolls, Network: Computation in Neural Systems **2**, 371 (1991).
[2] A. Treves, Physical Review A **42**, 2418 (1990).
[3] A. Treves, S. Panzeri, E. T. Rolls, M. Booth, and E. A. Wakeman, Neural Computation **11**, 601 (1999).

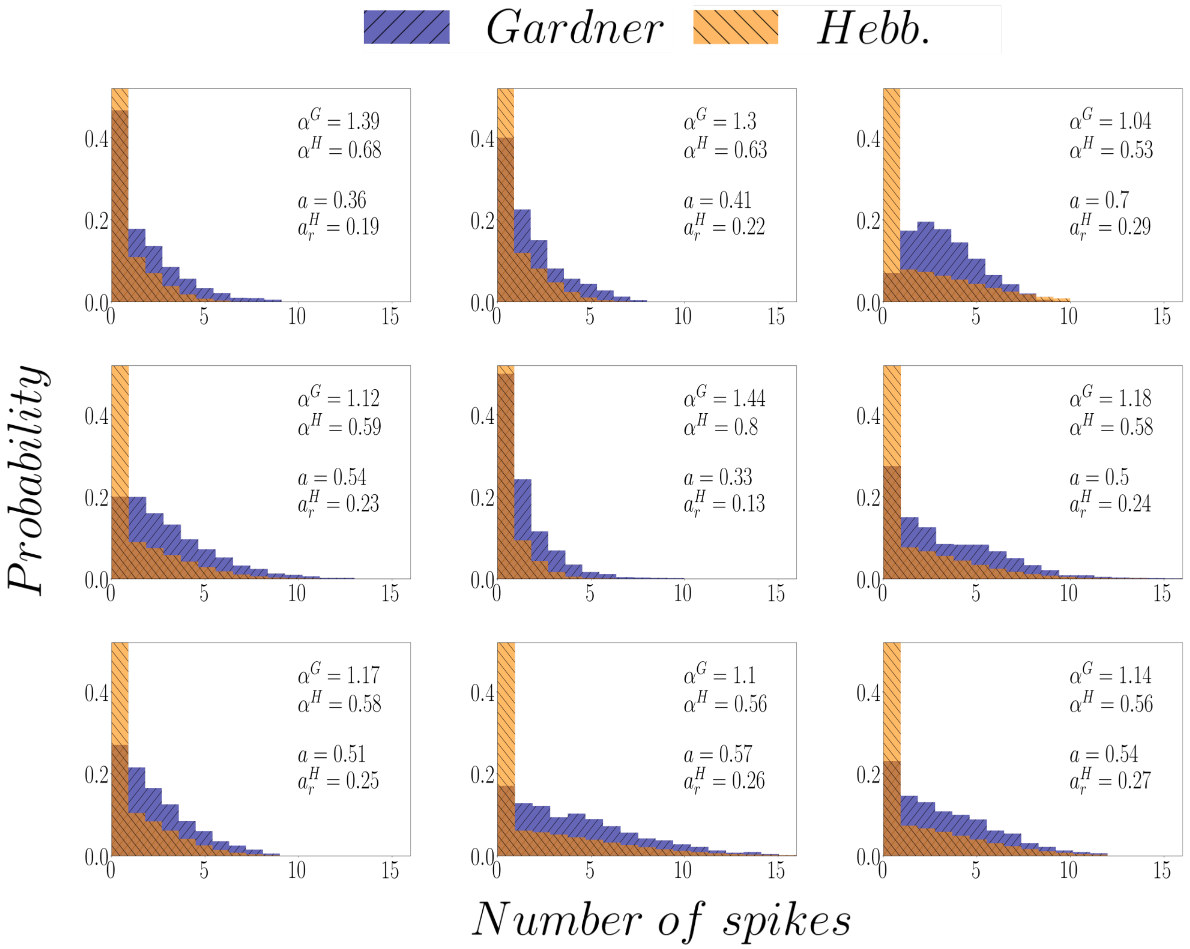


FIG. 2. Suplementary to Fig (3).

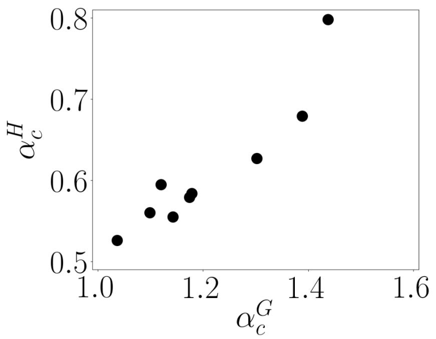


FIG. 3. Comparison between the values of the storage capacity *à la Gardner* and Hebbian, for the 9 empirical distributions extracted from [3].
